# gtexture: novel extension of image texture analysis to graphs and its application to cancer informatics

**DOI:** 10.1101/2022.11.21.517417

**Authors:** Rowan J Barker-Clarke, Davis Weaver, Jacob G Scott

## Abstract

**Objective:** The calculation of texture features, such as those derived by Haralick *et al*., has been traditionally limited to 2D-imaging data. We present the novel derivation of an extension to these texture features that can be applied to graphs and networks and set out to illustrate the potential of these metrics for use in cancer informatics.

**Approach:** We extend the pixel-based calculation of texture and generate analogous novel metrics for graphs and networks. The graph structures in question must have ordered or continuous node weights/attributes. To demonstrate the utility of these metrics in cancer biology, we demonstrate these metrics can distinguish different fitness landscapes, gene co-expression and regulatory networks, and protein interaction networks with both simulated and publicly available experimental gene expression data.

**Main Results:** We demonstrate that texture features are informative of graph structure and analyse their sensitivity to discretization parameters and node label noise. We demonstrate that graph texture varies across multiple network types including fitness landscapes and large protein interaction networks with experimental expression data. We show the ability of these texture metrics, calculated on specific protein interaction subnetworks, to classify cell line expression by lineage, generating classifiers with 82% and 89% accuracy.

**Significance:** Graph texture features are a novel second order graph metric that can distinguish cancer types and topologies of evolutionary landscapes. It appears that no similar metrics currently exist and thus we open up the potential derivation of more metrics for the classification and analysis of network-structured data. This may be particularly useful in the complex setting of cancer, where large graph and network structures underlie the omics data generated. Network-based data underlies drug discovery, drug response prediction and single-cell dynamics and thus these metrics provide an additional tool in tackling these problems in cancer.

## Introduction

Topology and texture are intrinsic properties of biological data and have been studied directly and indirectly across biomedical research. Textural and topological analysis methods have, for example, defined solutions in medical imaging classification, the analysis of biological signaling networks and modules, and in understanding the structure genotype-phenotype maps of evolution (**Fig. 1**). These textures and topologies have been used to identify biologically meaningful structures or patterns and in some applications to cancer, even been associated with clinical outcomes(Somasundaram, Litzler, et al. 2021).

**Figure 1.**
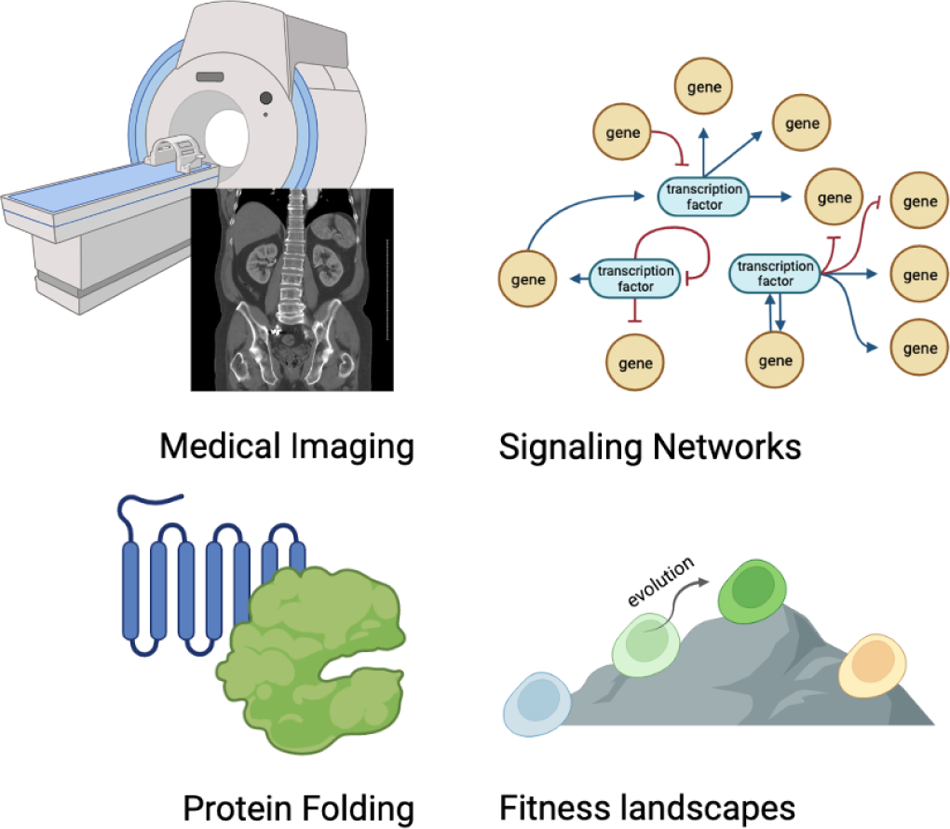
Illustration of areas in which topology is studied in biomedical research. Textural and topological studies are carried out in medical imaging, protein folding, signaling network and fitness landscape analysis.

Image“texture” features are a staple within image classification, with the gray-level co-occurrence matrix (GLCM) and associated statistics having been popular for decades(Haralick, Shanmugam, and Dinstein 1973; Haralick 1979). GLCMs are 2D histograms that record the frequency of neighboring pixel gray-level values in an image. The Haralick texture features are second-order statistics that summarize the distribution over pixel-neighbour pairs. These statistics include measures that reflect heterogeneity, homogeneity, and contrast between neighboring pixels within images. These are very commonly used in medical physics where texture features from CT and MRI images have been related to tumor type, severity, and prognosis (Mohanty, Beberta, and Lenka 2011; Yang et al. 2012; Zulpe and Pawar 2012; Jain 2013; Torheim et al. 2014; Novitasari et al. 2019). We note that co-occurrence matrices, although most commonly used in imaging, have also been extended to NLP fields (Momtazi, Khudanpur, and Klakow 2010; Benoit et al. 2018), audio processing (Terzopoulos 1985; Sayedelahl et al. 2011; Muhammad et al. 2017) and recently in pathology in a form derived by Saito et. al., describing the co-occurrence of nuclear features in physical cell neighborhoods (Saito et al. 2016).

Recent interdisciplinary work has successfully extended different graph-based topological analyses to image-derived point clouds and more recently to images themselves, including the use of cubical complexes to derive prognostic topological features from medical images (Lawson et al. 2019; Hajij, Zamzmi, and Batayneh 2021; Somasundaram, Litzler, et al. 2021; Somasundaram, Wadhwa, et al. 2022).

Topology has also shed light on biological networks. As increasing amounts of proteomic and transcriptomic data become available, a wealth of information about gene expression and protein-protein interaction networks arises. Within cancer, the dysregulation of signaling pathways and modified interactions between mutant proteins means that holistic network analyses may potentially identify critical features in these data sets. Topological analysis of gene and protein networks has identified regulating gene sub-networks for potential drug targeting, improved understanding of the stability of gene signaling networks, and even given prognostic indications in breast cancer (Sardiu et al. 2019; Kumar, Blondel, and Extavour 2020; Guo and Amir 2021; Yin et al. 2021; Weaver, Pishas, et al. 2021).

Another area within biology in which topology has been of interest is the study of fitness landscapes(Lum et al. 2013), a particular subclass of networks. Fitness landscapes typically encode a genotype space and associated fitness. In cancer, these are of particular interest as fitness landscapes encode the constraints of Darwinian evolution and are informative in the modeling of resistance and optimization of treatment (J. Scott and Marusyk 2017; Nichol, Rutter, et al. 2019; King et al. 2022). As the topology of a landscape can restrict or promote access to certain evolutionary trajectories, constraining the accessibility of local and global maxima (Levinthal 1997) measures have been developed to evaluate landscape “ruggedness” (Barnett et al. 1998). Modeling of “tunably rugged” landscapes has allowed the direct exploration of the effect of topology and texture upon evolution, demonstrating strong associations with evolutionary timescales and outcomes (Kauffman and Weinberger 1989; Barnett et al. 1998; Franke et al. 2011). As the ability to engineer and measure fitness landscapes experimentally has become easier, the nature of fitness landscapes is of growing interest; particularly in modern studies of evolutionary cancer therapies, drug resistance, and biological control(Nichol, Jeavons, et al. 2015; Diaz-Uriarte 2018; Nichol, Rutter, et al. 2019; Hosseini et al. 2019; Iram et al. 2021; Hsu et al. 2022).

The aim of this work is to extend topological research by bringing the tools of image analysis to analyze network structures. In the case of networks with node labels, the underlying network can be considered a fixed wiring diagram that connects node values. In the case where only the node labels are dynamic, typical graph summary measures such as the number of nodes, number of edges, maximum degree, minimum degree, average degree, diameter, average path length, and edge density remain unaffected by varying node weights. These new Haralick-derived graph texture metrics allow for the extraction of second-order features influenced by topology and node value, allowing for comparative analysis of data with accompanying knowledge networks. In cancer contexts these node labels may represent expression, growth rates or frequencies, that may vary across time in evolutionary contexts.

While this extension to graphs is novel in itself, we focus on the use and potential of these GLCM-equivalents and Haralick texture features in cancer biology and calculate them for several biological network types. We analyze networks with accompanying categorical and continuous node attributes, demonstrating this method on examples of idealized artificial gene regulatory networks, evolutionary fitness landscapes, and human protein-interaction networks with publicly available experimentally derived cancer cell line expression data.

Our R package for the calculation of these graph texture features, **gtexture**, is available on github at [removed for double-blind]

## Methodology

We show how co-occurrence matrices and texture calculations can be generated from and applied to graph objects. Co-occurrence matrices are 2D histograms, traditionally reflecting the pairwise distribution of neighboring pixel values in images. To apply this method to graphs or networks they must have node attributes or weights. These weights can be in the form of discrete weights or ordered categorical attributes. Broadly speaking we extend image texture metrics to graphs by considering node attributes to be analogous to pixel values and a node’s edges to be equivalent to pixel neighborhoods.

### Graph definitions

A graph *G* can be defined as a pair (*V, E*) where *V* is a set of vertices representing the nodes and *E* is a set of edges representing the connections between the nodes. We define the set of edges *E* as, *E* = (*i, j*)*|i, j ∈ V* where each edge is the single connection between nodes *i* and *j*. In this case, we say that nodes *i* and *j* are neighbors. For our current versions of these metrics, the edges of the graph must be unweighted and each node must have a node weight or ordered category, *w_i_*, associated with it, where *i ∈ V*. Co-occurrence matrices can be described in network terms as node-weight adjacency matrices.

Whilst we believe there are no directly comparable second-order graph metrics(Li et al. 2012), we utilize a few existing metrics and summary statistics of node-weighted graphs for comparison. The outline of the method and approach underlying the discretization, co-occurrence, and texture calculation for our metrics follows below.

### Discretization

Given a number of nodes *n*, a network’s adjacency matrix is size *n × n*. If the number of distinct node weights is *w*, the dimension of the co-occurrence matrix, *C*, is *w×w*. Co-occurrence matrices summarize a network when the number of distinct node weights is less than the number of nodes, *w < n*. Although this is already the case for some networks, we provide methods to reduce the number of unique node weights, including node weight binning options for continuous node weights within the package. Continuous data can be transformed via several discretization methods.

The following methods of discretization can be found within the package and two of them are demonstrated within **Figure 2**.

**Figure 2.**
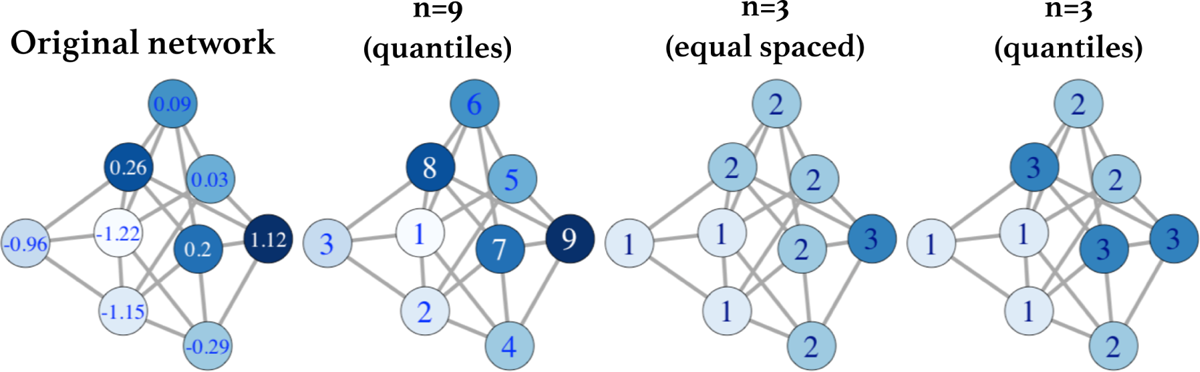
A demonstration of different discretization methods for continuous node values is shown. One example of a randomly generated undirected network with different random continuous expression values attributed to the nodes is shown. Discretization with 9 quantile levels matching the number of unique values and 3 levels with both equally spaced numerical bins and with 3 levels assigned to tertile groups are shown.

**Equal:** We can use a breaks method to slice the node weights into *n* equally spaced levels containing potentially different proportions of the data.

**Quantiles:** In this method the values are split into *n* groups containing equal numbers of values.

**k-means:** Values are split into *n* = *k* groups using 1D kmeans clustering.

### Co-occurrence matrix calculation

For any graphical structure, the edges between nodes are captured in an adjacency matrix. These edges are used for the calculation of the distribution of co-occurring neighbor pairs. In an undirected network (symmetric adjacency matrix), the neighboring node values are summed over all edges. In a directed graph, the adjacency matrix is used directly to iterate through pairs of connected node values in a single direction. The element *C_i j_* of the co-occurrence matrix is the number of times within the network a node with weight *i* shares an edge with a node of weight *j*. Examples of two separate co-occurrence matrices for a toy gene regulation network with four bins of expression values are shown in **Fig. S1.**

### Graph texture metric definitions and interpretation

Standard image analysis practice uses the co-occurrence matrix to generate texture features for the image. Haralick defined several statistical features and these calculations on the co-occurrence matrix traditionally reflect properties of an image’s texture (Haralick, Shanmugam, and Dinstein 1973; Haralick 1979). The mathematical definitions of the eleven key texture features calculated in this paper are shown in **Table 1**. Although the direct mapping of image texture features to visible texture changes is not fully understood, we discuss the analogous interpretations based on the definitions of the features in the graph texture metric setting. Our package extracts these features and in order to compare these features across different categories of network, metrics are normalized across compared groups.

**Table 1.**
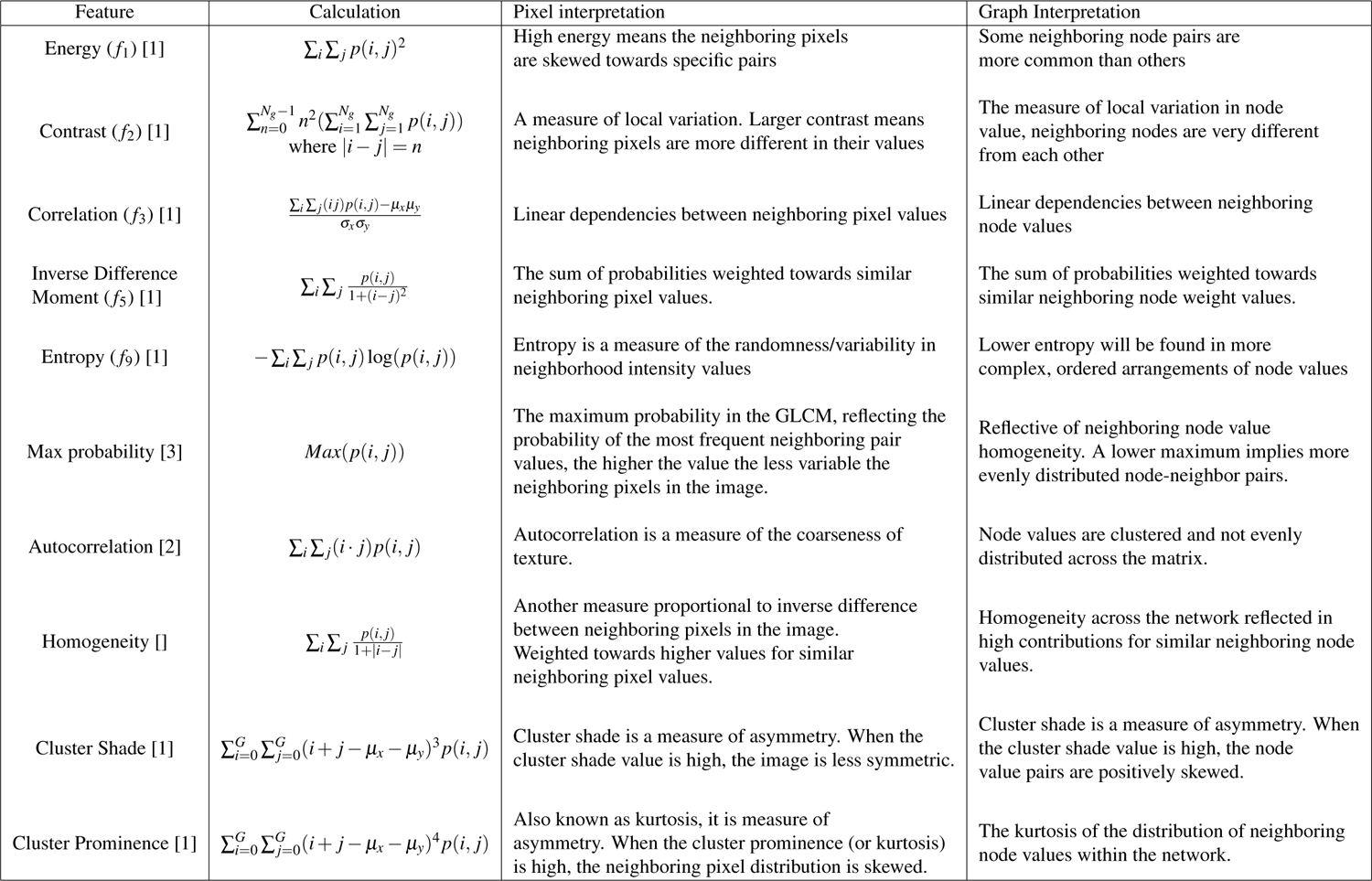
Definitions of several original Haralick texture features and other key GLCM statistics. Interpretations in the image context and the analogous interpretations in the graph or network context are shown. [1](Haralick, Shanmugam, and Dinstein 1973) [2](Soh and Tsatsoulis 1999) [3](Clausi 2002)

### Network examples

To demonstrate the generation and meaning of these metrics we used multiple biologically inspired network examples. We utilize constructed toy networks, informed gene expression networks, and fitness landscapes.

We constructed three different examples of biological networks with a small number of gene modules (n=3 or 5) and modularity scores of 0.5-0.7. We used the code from work by Sah et. al. to generate examples of modular graphs (Sah et al. 2014). To examine the sensitivity of the metrics we compared these constructed ordered and modular networks with specific node values to the same networks with bootstrapped node weights and to metrics on the same networks with added noise.

Another specialized network type is the evolutionary fitness landscape. Genotypes in the fitness landscape are neighbors, connected by an edge if they are accessible through a single evolutionary timestep (eg. mutation). The underlying network structure is defined by this evolutionary access and the node weights are the fitness values. As the number of available experimental fitness landscapes is limited, we used statistically generated landscapes generated via the packages *fitscape* and *OncoSimulR*.

We utilized basic landscape networks with specific fitness distributions to demonstrate our methodology, explain what the metrics summarize, and begin to probe the potential in cancer biology for these metrics. Utilizing the R package **OncoSimulR** (Diaz-Uriarte 2017) we generated three classes of basic model landscape and sets of NK landscapes and converted these into fitness landscape objects using the R package *fitscape*. The *OncoSimulR* package utilizes MAGELLAN, a fitness landscape analysis toolset, (Brouillet et al. 2015) to generate some standard models of fitness landscapes; additive, eggbox, and house of cards.

**Additive model landscapes** In the additive model, mutations have a specific fitness increase or decrease and multiple mutations increase or decrease fitness in a linear, additive fashion. These landscapes are very smooth and monotonic.

**Eggbox model landscapes** In the eggbox model there are only 2 different possible fitness values, the base fitness and base fitness + *e* (the “height” of the eggbox), thus any mutation moves a genotype from low to high fitness or vice-versa, and neighboring genotype fitness values are always distinct.

**House of cards model landscapes** The House of Cards (HOC) model is a name for a random fitness model, here the fitnesses of different genotypes are uncorrelated and not dependent on the genotype, this is an effective null/random model.

For each basic network type, we generated a set of 4 allele, 16 genotype fitness landscapes for analysis. We created sets of ten random additive, eggbox, and House of Card landscapes.

We also simulated “NK” landscapes with the same package, comprising 5 alleles (32 genotypes) and varying the epistatic interaction K from 1 to 3. For each value of K (1 to 3), we generated 500 random “NK” landscapes. We compared these to some traditional measures (roughness:slope ratio) of landscape ruggedness.

### Simulated gene expression

In order to analyze gene expression within graphical structures of established human protein-protein interaction networks, we used STRINGDB to obtain pathway-specific subnetworks, the KEGG database, and the *KEGGGraph* package to convert between gene and protein identifiers.(Ogata et al. 1998; Zhang and Wiemann 2009; Szklarczyk et al. 2015). To use experimentally derived gene expression as node values for these networks we used the publicly available Cancer Cell Line Encyclopedia (CCLE) gene expression dataset (Barretina et al. 2012). We also used the R package *graphsim* as a method of simulating gene expression values on PI3-Kinase and TGF-*β* co-expression networks with varying correlation strength(Kelly and Black 2020). In order to evaluate our metrics upon simulated gene expression, we varied the correlation parameter of the gene expression simulation from 0.2 to 0.8. Expression levels from both the simulation and the CCLE data were discretized into 4 node levels and these expression values were used as node weights.

### Subgraph generation

In order to identify relevant subnetworks of the human protein-protein interaction network we used the R package *crosstalkr* to identify subnetworks based on proteins of interest and their interactions(Weaver and J. G. Scott 2023). The crosstalk between a set of proteins of interest was generated based on random walks of length 100 and minimum connectivity score for edges of 1.

### Random forest classifiers

To train our classifiers we split the 1406 cell-lines from the CCLE dataset at random into 70:30 training and test split. We used the R package *randomForest* to build random forest classifiers. This package generates random forest classifiers using Breiman’s random forest algorithm. Our classifiers were built on the training sets using a generation of *n* = 500 trees per classifier.

## Results

In order to demonstrate both the efficacy and potential of texture analysis as applied to networks we apply our method to a selection of biological and cancer-specific networks. Networks and graphs, as a general mathematical structure, have been great tools for encapsulating many types of biological information with underlying network structures. To reflect this variety we utilize several categories of graphs in order to demonstrate co-occurrence calculation and texture feature generation. Due to the novelty of these graph metrics and the intrinsic heterogeneity and complexity of experimental biology, we include examples of artificial modular gene networks and idealized model fitness landscapes before assessing the use of these metrics on knowledge-based gene and protein interaction and expression networks. We compare metrics using a range of simulated, noisy, and experimentally derived publicly available gene expression data representing the node labels.

### Metrics reflect differences in artificial biological networks

In order to demonstrate and assess these novel metrics, it is important at first to examine the metrics on networks with known features and properties. We constructed three examples of artificial biological-type networks (Sah et al. 2014). We allocated the nodes representing genes in each community cluster on the network with the same node value.

These toy networks also allow us to further explain these Haralick metrics in the graph context. We see (**Fig. 3**)that in each case (A, B & C) that the original graphs are highly organized, and thus the neighboring node value pairs in the network are asymmetric, reflecting higher probabilities of certain types of neighbor node-label pairings in each case. For example in graph (A) none of the clusters are ever connected to neighbors of the same color/cluster, this gives this network a high energy and low entropy but also very low homogeneity as compared to the other graphs. In graphs (B) and (C), the connectivity between genes within clusters gives these networks a high modularity and their low connectivity to the rest of the network gives these networks a high homogeneity and autocorrelation compared to randomly allocated node values. We can see that in (C), the lower number of clusters or levels for the node values equates to a higher max probability value than in (B).

**Figure 3.**
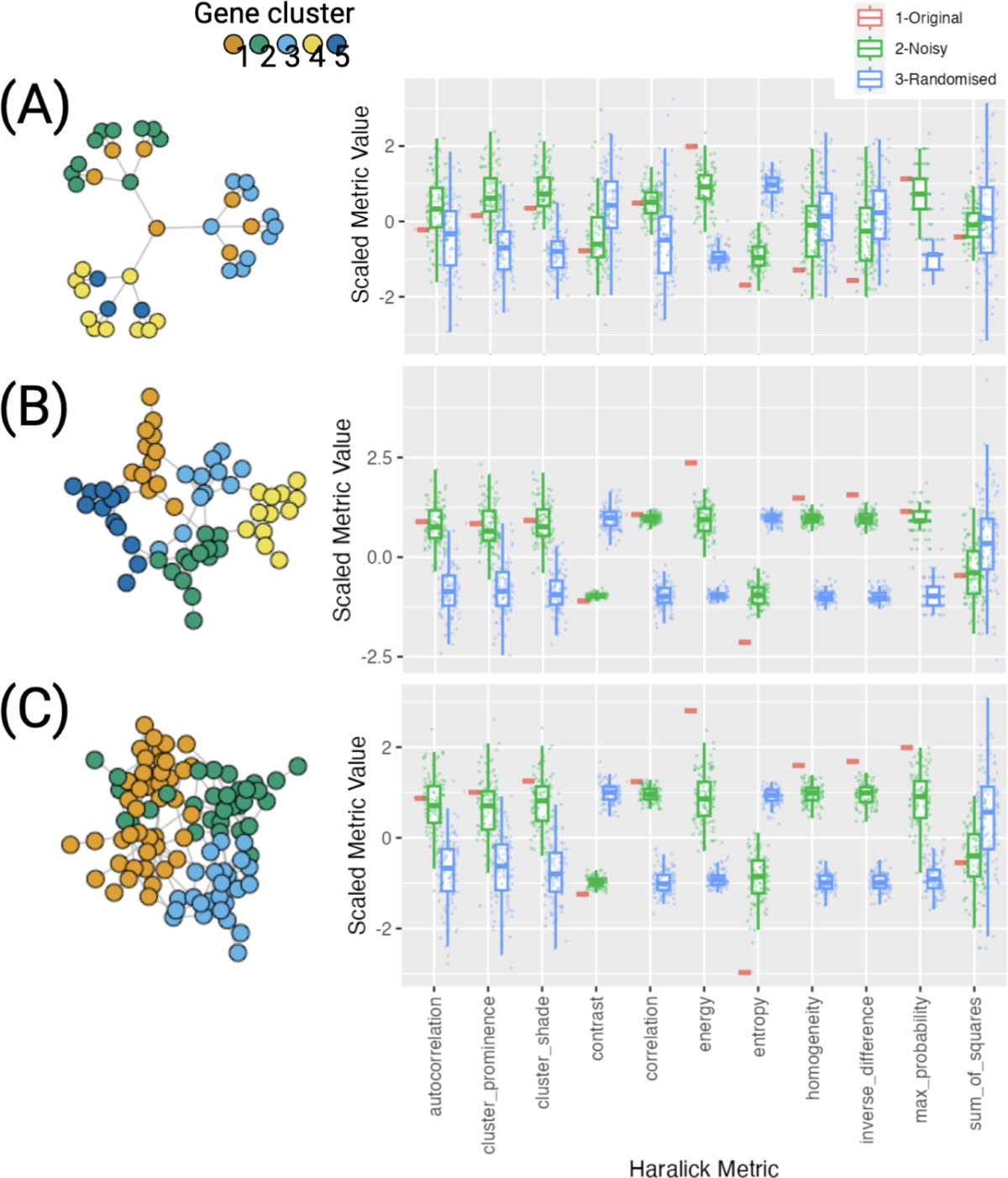
Three examples of simple possible biological gene networks. Texture features were generated with original graphs (red) and clusters were given equal node values for expression. We added noise (green) in addition to comparing metrics for randomly sampled node values(blue). These simple graph structures help to illustrate the texture metrics in their non-image form.

### Metrics are sensitive to noise and number of discrete node levels

Using an initial example of a modular graph (Q=0.7), generated with five node values assigned to five module clusters, we calculated the Haralick graph metrics for this graph, varying the strength of noise and the number of discrete levels into which to bin node values(**Fig. 4**).

**Figure 4.**
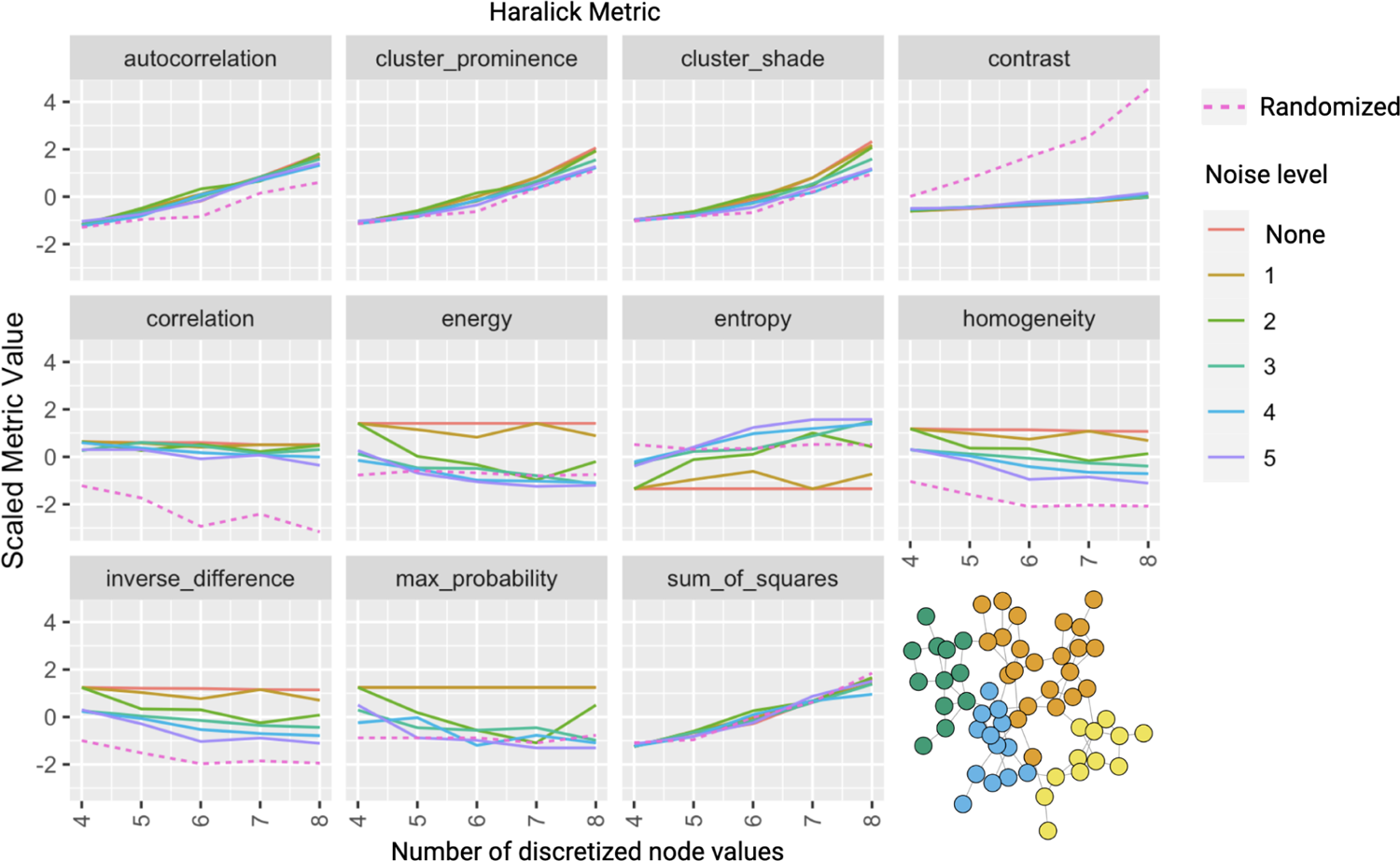
Scaled Haralick feature values vary with noise and with number of discrete node levels. One example of a randomly generated modular network with four initial node values is used and both the number of discrete levels (*x*-axis) and magnitude of added uniform noise(color) are varied with the effect on each metric shown.

Traditional Haralick features are sensitive to the number of gray levels chosen for the co-occurrence calculation(Löfstedt et al. 2019) and to noise(Brynolfsson et al. 2017; Schad 2022). We expect similar sensitivity in our metrics due to their analogous form.

Varying both noise and number of bins for the discretization of the node weights changes feature values (**Fig. 4**). We utilize randomization of the node values for comparison. For this network example, the correlation and contrast are relatively robust to noise and the number of levels in comparison to a randomized node assignment. In comparison, the sum of squares and cluster prominence are highly sensitive to both.

### Metrics reflect topologies of fitness landscapes

Fitness landscapes can be considered a special case of a biological graph, the wiring diagram or edges in the graph can be defined by mutation and the node values encode the fitness (often represented by the growth rate) of each of the underlying genotypes. The distribution of fitness values and their connectivity (ie topology and texture) of a genotype landscape are associated with evolvability as the landscape restricts movement towards fitter genotypes via mutation and selection.

Co-occurrence matrices and graph texture metrics may valuable information generated from a fitness landscape, as these encode properties of the distribution of neighboring fitness values. In order to assess whether our Haralick graph features are meaningful metrics for fitness landscapes we compared three distinct idealistic landscape types and a commonly used set of tunably rugged landscapes. Illustrative examples of these landscapes are shown (**Fig. 5a**). We tested our pipeline on these models using 4-level node weight equal discretization on 4 alleles (16 genotypes) model landscapes.

**Figure 5.**
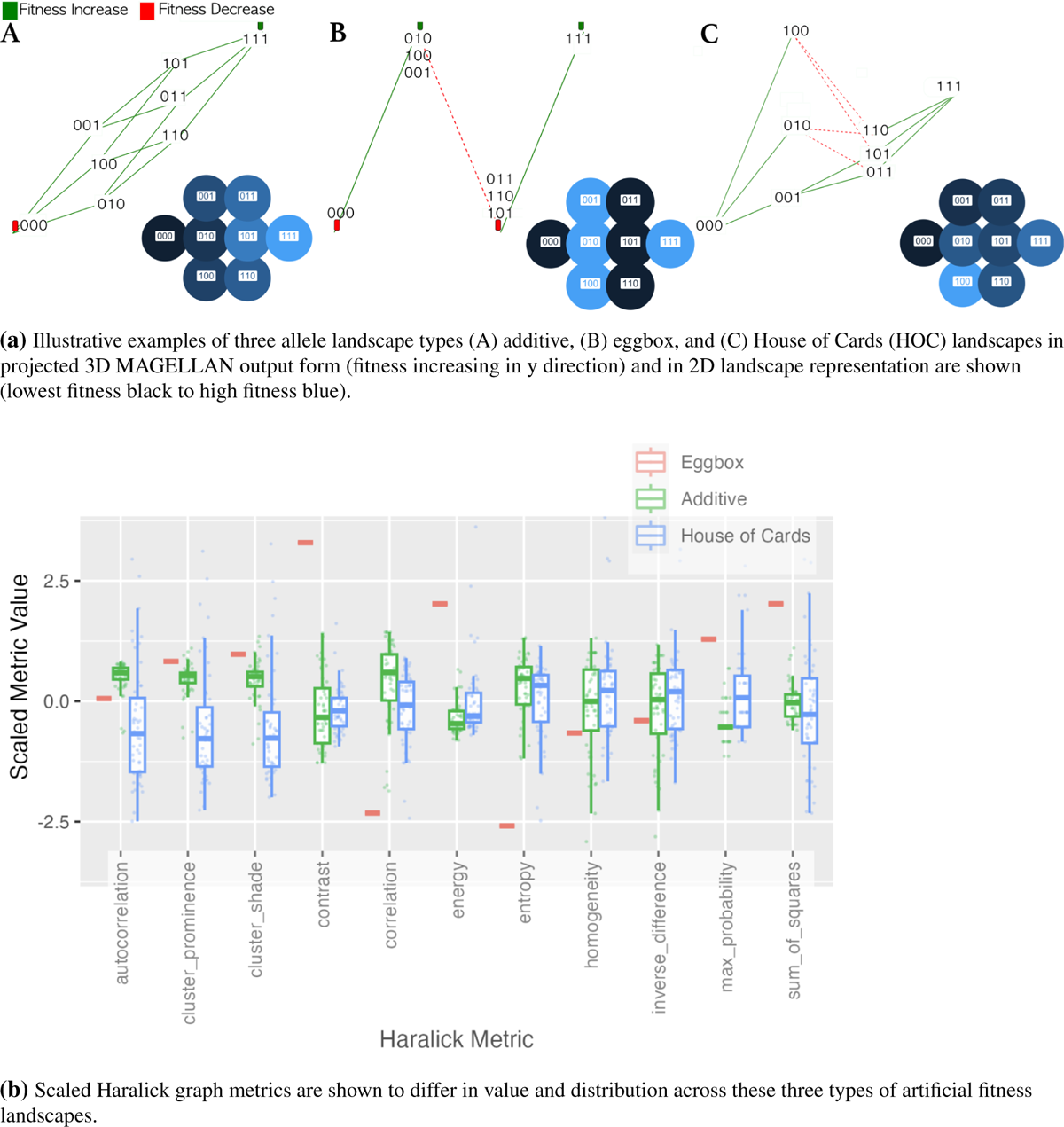
Illustration of landscapes and distribution of GLCM metrics on them. **a)** Illustrative landscapes are shown for each type. **b)** “Eggbox” landscapes collapse under discretization and normalization. The eggboxes have the highest contrast and lowest homogeneity as neighboring genotypes have alternating fitnesses, the additive model shows the highest correlation, homogeneity, and lowest contrast whereas the house of cards (HOC) model with its random fitnesses shows the largest range of values due to a wider spread of neighboring fitness pairs.

The Haralick texture features are calculated on these landscapes, and the normalized metrics are shown in **Figure 5b** for comparison. The results and distributions of these features reflect our understanding of the metric definitions. The eggbox landscapes show extremely high contrast and low entropy due to the alternating nature of the landscape, where the fitness of neighboring genotypes always alternates between peaks and valleys of equal height and depth respectively. The eggbox landscape also shows relatively high cluster prominence and shade and low homogeneity for the same reasons. As expected the additive landscape shows high neighbor correlation and highest neighbor autocorrelation, reflecting the smooth and monotonically increasing fitness surface. Meanwhile, the random landscape (“House of Cards”) shows the largest variation across all metrics and the lowest autocorrelation as would be expected from random fitness assignments.

Whilst the cancer fitness landscapes measured will be undoubtedly more complex than the model landscapes above there exist a standard set of landscapes used for evolutionary computation that have more similarities to experimental landscapes(Kauffman and Weinberger 1989; Wang and Dai 2019). We included an analysis of the texture of simulated sets of these tunably rugged “NK” landscapes (Fig. S2). As epistatic interactions increase, the contrast between neighboring fitness values decreases (therefore dissimilarity decreases). At K=0, the landscape is smooth and additive. As K is increased, the landscape becomes more rugged as epistatic interactions increase, and the correlation increases with K.

### Texture metrics vary between experimental and simulated gene expression data on the same network

The development of biological networks has been driven by growing works in transcriptomic and proteomic studies. Protein-protein interaction (PPI) networks have been built based on experimental evidence probing the interaction of different proteins. We hypothesized that our technique may be useful in assessing experimental data gathered in different samples for established biological networks, in particular as a way of summarizing expression patterns across different topologies of protein interaction networks. We examined the phosphoinositide-3-kinase (PI3K) cascade network and assigned gene expression values to nodes, using both the CCLE experimental dataset and simulated through the *graphsim* package.

**Figure 6** shows expression on the PI3K network and how the Haralick metrics vary with increasing expression correlation. In the network describing PI3K regulation we see the expected results, that contrast decreases, correlation increases, entropy decreases and homogeneity increases as correlation in the underlying expression simulation increases. When the gene expression data from the cancer cell lines in the CCLE is compared, we see that these are significantly different (more extreme) than metrics upon the simulated expression, showing results that correspond to increased correlation strengths, lying outside the simulated distributions. These metrics may provide a way to assess and improve methods for simulating gene expression.

**Figure 6.**
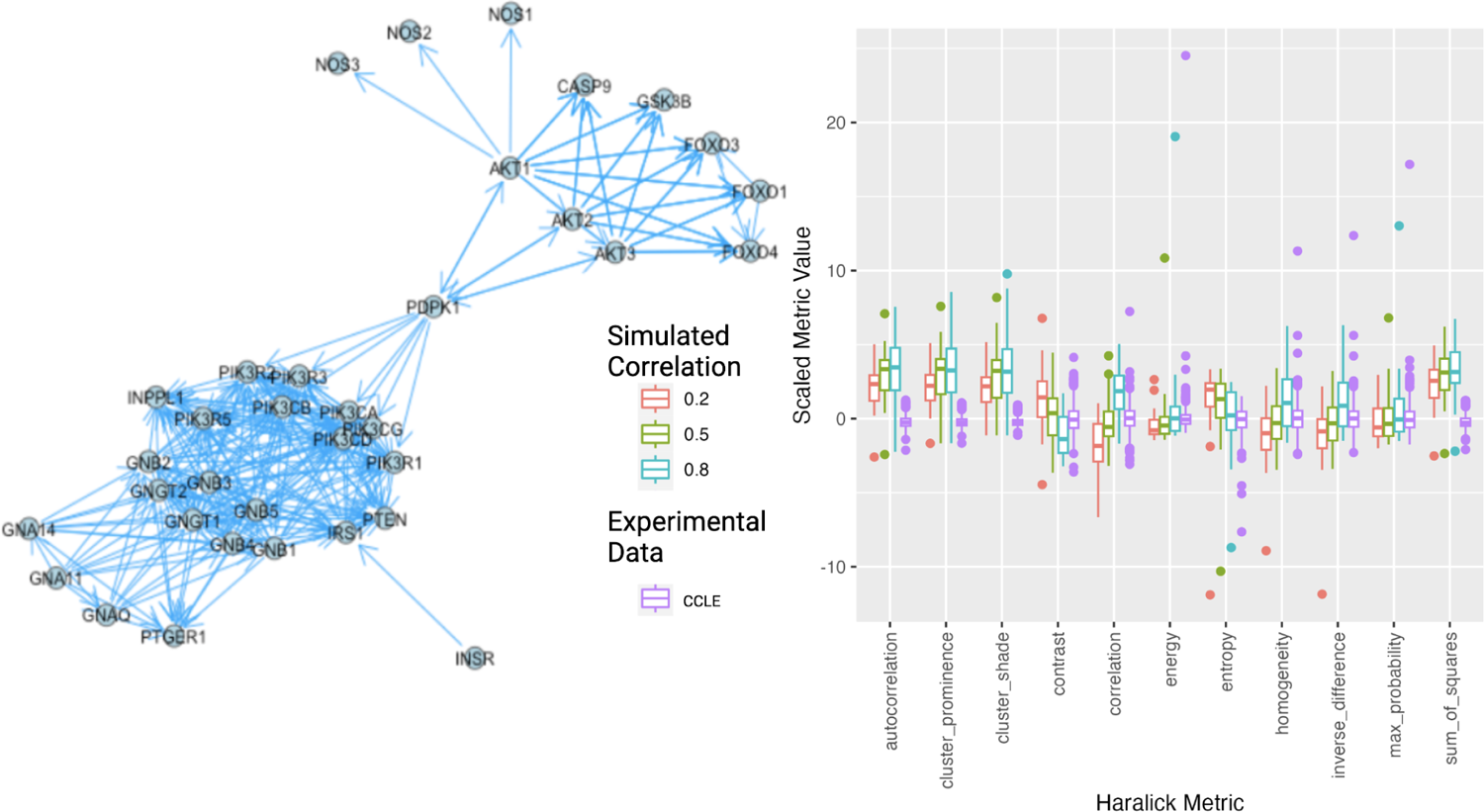
Phosphatidylinositol-3-kinase interaction network. Texture features were generated with simulated expression data and experimental CCLE gene expression data (pink). Plots of graph texture features are shown for simulated gene expression on the PI3K gene network with different strengths of co-expression correlation show a trend across node value correlation strength.

### Metrics can be used to classify cancer lineages from expression networks

In order to better address the utility of these metrics within cancer we decided to examine whether the metrics could be used for classification of CCLE data. We decided to use the metrics to classify expression patterns in biological sub-networks, we identified subnetworks with expected differences in regulation and expression patterns between cancer cell lines of varying cell lineage. We calculated the metrics for expression on the EGFR signaling pathway subnetwork. We show the distribution of our texture metrics on this network for some of the most common cancer subtypes within the CCLE dataset. EGFR (epithelial growth factor) dysregulation is associated with solid tumors and we see corresponding differences in metric values between the epithelial (solid - lung, breast, ovary, central nervous system, prostate, and skin) and non-epithelial (blood, lymphocyte) cell line samples (**Fig.** S3). EGFR amplification is a particularly common feature of glioblastoma, a large proportion of CNS tumors, and we see this reflected in more extreme metric values for CNS tumors. To assess whether there are differences between the metrics for primary and metastatic samples in tumors with likely EGFR dysregulation, we also analyzed the same metrics between primary and metastatic cell lines with central nervous system origin (**Fig.** S4). We find significant differences in the metric distributions between primary and metastatic cell lines.

To demonstrate how these metrics can potentially be used for classifying graph-structured biological data we created a random forest classifier using our graph texture metrics. We tested two classifiers, the first aimed to classify cell line expression data for lung and blood lineage cell lines, the two largest proportions of the data set(*n* = 300). We built a random forest classifier on a training subset of these cell lines and validated it on the test set. Our model performs with an accuracy of 89.7%. The confusion matrix (**Table 2a**) and confidence intervals are shown. The second classifier used a subgraph of interactions between the expression of genes conserved across epithelial-mesenchymal transitions (Vimentin, EPCAM)(Cook and Vanderhyden 2020). We analyzed the ability of this classifier to distinguish central nervous system and lung cancer cell-lines. This classifier performed with an accuracy of 82.1%.

**Table 2.**
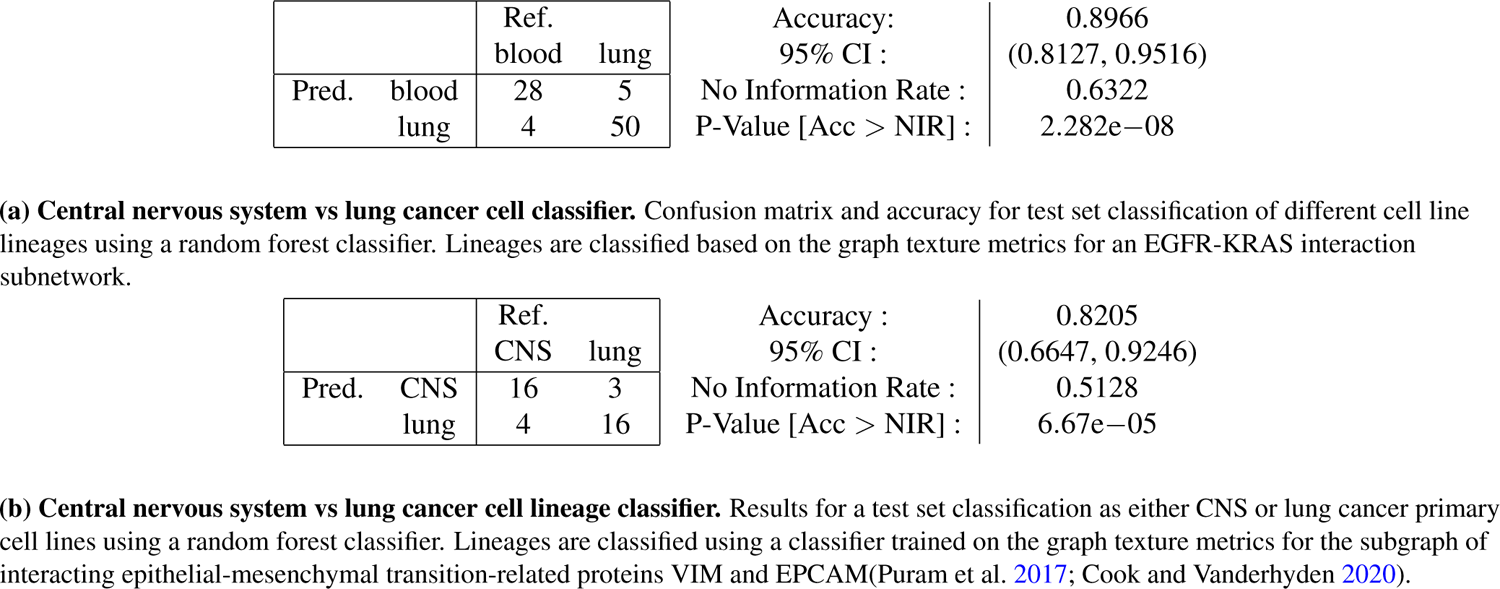
Confusion matrices and classifier accuracies for texture based random forest lineage classifiers.

|  |  |  |  |  |  |
| --- | --- | --- | --- | --- | --- |
|  |  | Ref. |  | Accuracy: | 0.8966 |
|  |  | blood | lung | 95% CI : | (0.8127, 0.9516) |
| Pred. | blood | 28 | 5 | No Information Rate : | 0.6322 |
|  | lung | 4 | 50 | P-Value [Acc > NIR] : | 2.282e−08 |
|  |  | Ref. |  | Accuracy : | 0.8205 |
|  |  | CNS | lung | 95% CI : | (0.6647, 0.9246) |
| Pred. | CNS | 16 | 3 | No Information Rate : | 0.5128 |
|  | lung | 4 | 16 | P-Value [Acc > NIR] : | 6.67e−05 |

(b) **Central nervous system vs lung cancer cell lineage classifier.** Results for a test set classification as either CNS or lung cancer primary cell lines using a random forest classifier. Lineages are classified using a classifier trained on the graph texture metrics for the subgraph of interacting epithelial-mesenchymal transition-related proteins VIM and EPCAM(Puram et al. 2017; Cook and Vanderhyden 2020).

## Discussion

Cancer informatics is rich with graph-structured data, between knowledge graphs for drug discovery and protein and gene expression datasets generated in bulk and single cell experiments, these networks provide a rich resource for novel analysis methods. With the expanding prevalence of networks with structure and variable node labels, there exists a need for cross-disciplinary analysis methods aimed at analysing networks and graphs that extend beyond their topology alone. Network analysis techniques and summary statistics typically assess edge properties and topology, but experiments contain large amounts of additional data about the nodes of a network, for example, a gene or protein, or cell-line. In order to analyze these node properties in tandem with the network, we demonstrate, for the first time, the generation of co-occurrence matrices and graph texture features as summary features of general networks.

Co-occurrence matrices upon networks reflect the distribution of connected pairs of node values within a network or graph object. Specific examples include the gene expression of neighboring genes in a network or fitness values of neighboring genotypes in a fitness landscape. Suitable networks for this metric must have ordered node attributes or discrete or continuous node weights.

Our results demonstrate stark differences in texture between network types toy gene networks, cancer gene expression networks, and simulated fitness landscape networks. As this is a new methodology,we initially present these metrics upon interpretable and well-understood network examples. We use these to demonstrate that like equivalent image texture features, these metrics are sensitive to node label discretization and also to noise. Some features are more robust than others and just as in image classification, these sensitivities should be considered when applying these methods. Future work could derive invariant versions of these metrics in analogous ways to the recent gray-level invariant texture features for images(Löfstedt et al. 2019). Our method showed that the graph texture features calculated on different landscapes and networks of the same size but with different topologies vary. We demonstrate that these features correspond to the properties of node neighborhoods and graph topological features. The method of calculating the distribution of neighboring pairs and subsequent statistics can therefore successfully be applied to networks with node attributes and can simultaneously measure the co-dependency of node labels and network topologies. The same methodology can be extended to calculate further novel network metrics derived from gray-level image metrics using similar principles. The package provides a framework for the future study of the optimization of parameters such as the number of discrete levels chosen to encode node values such as gene expression or fitness values. Although highly specific methods designed for detecting landscape ruggedness exist, this discretization and co-occurrence matrix method is more generalizable.

We also demonstrate differences in texture between cell lines when using experimental data from different cancer types. Although the metrics can distinguish certain cell-line lineages, the selection of appropriate subnetworks is still required to produce successful classification and further metrics including those reflecting topology node labels alone could be added to classification attempts as a way to potentially improve classification accuracy.

Although the GLCM texture features are well characterized in imaging, the true utility of these metrics upon networks has yet to be explored. By utilizing these ideas from image analysis, this method provides a simple analysis and summary technique that is particularly effective for larger network types with node-specific intensities. As the fitness landscape data generated and collected becomes larger, methods such as this that can reduce the dimensionality of complex networks while retaining information about the structure may be useful. As such, this package provides efficient computation of summary statistics for graphs with edges and discretization of node attributes.

We believe that this package can be applied to many network types, not just those represented here, and may be able to derive second-order statistics reflective of important network-node characteristics. This method can be applied, for example, to graphs used for literature searches and target identification, to fitness and growth rate data, gene expression, protein expression, time series data, and cross-sectional data. We encourage the use of this package in exploratory network analyses across cancer.

## Author’s contributions

## Supplementary Information

GLCM based texture features often comprise a subset or variation upon some core statistical metrics. Since the initial derivation of the Haralick texture features many additional features have been investigated. Some of the features that have been included in our package and can be considered upon networks include the following; Sum of Squares: Variance, Sum Average, Sum Variance, Sum Entropy, Difference Variance, Difference Entropy, Information Measure of Correlation 1 (IMC1), Information Measure of Correlation 2 (IMC2) and Max Correlation Coefficient (MAXC).

## Supplementary figures

**Figure S1.**
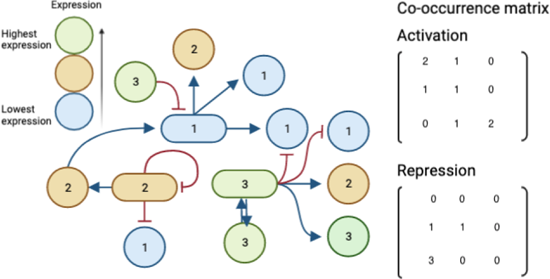
Co-occurrence matrices calculated on a toy gene regulation network. In the case of a directed graph only directions included are counted. In directed activation and repression graphs, two separate co-occurrence matrices can be calculated for the same network. When we vary the tunable ruggedness of simulated landscapes by varying K in the OncosimulR package, we see changes in the texture metrics (Supplementary Figure S2). Epithelial tumors are separated from non-epithelial tumors in the CCLE datasets. The EGFR network and associated expression data shows significantly different texture between primary and metastatic central nervous system tumors (Supplementary Figure S4)

**Figure S2.**
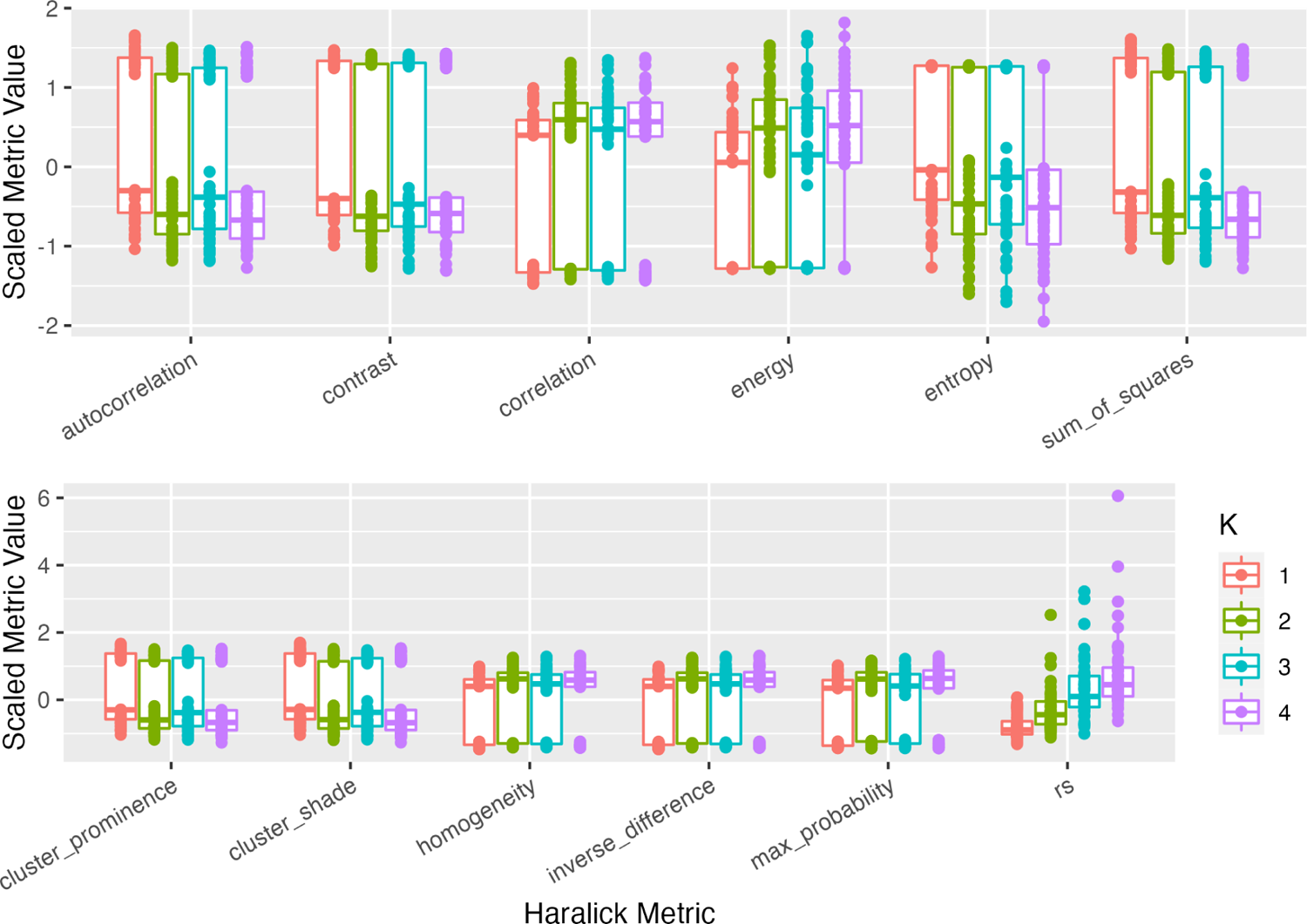
Metrics across NK landscapes. Haralick measures differ across different K for tunably rugged landscapes (5 alleles). Haralick features show distinct bimodal distributions of metrics for tunably rugged landscapes with fitnesses binned into 4 groups. The roughness-slope metric outperforms these in terms of separating landscapes with a single measure, but metrics contain information about landscape structure. Lines connect the same landscapes across different metrics.

**Figure S3.**
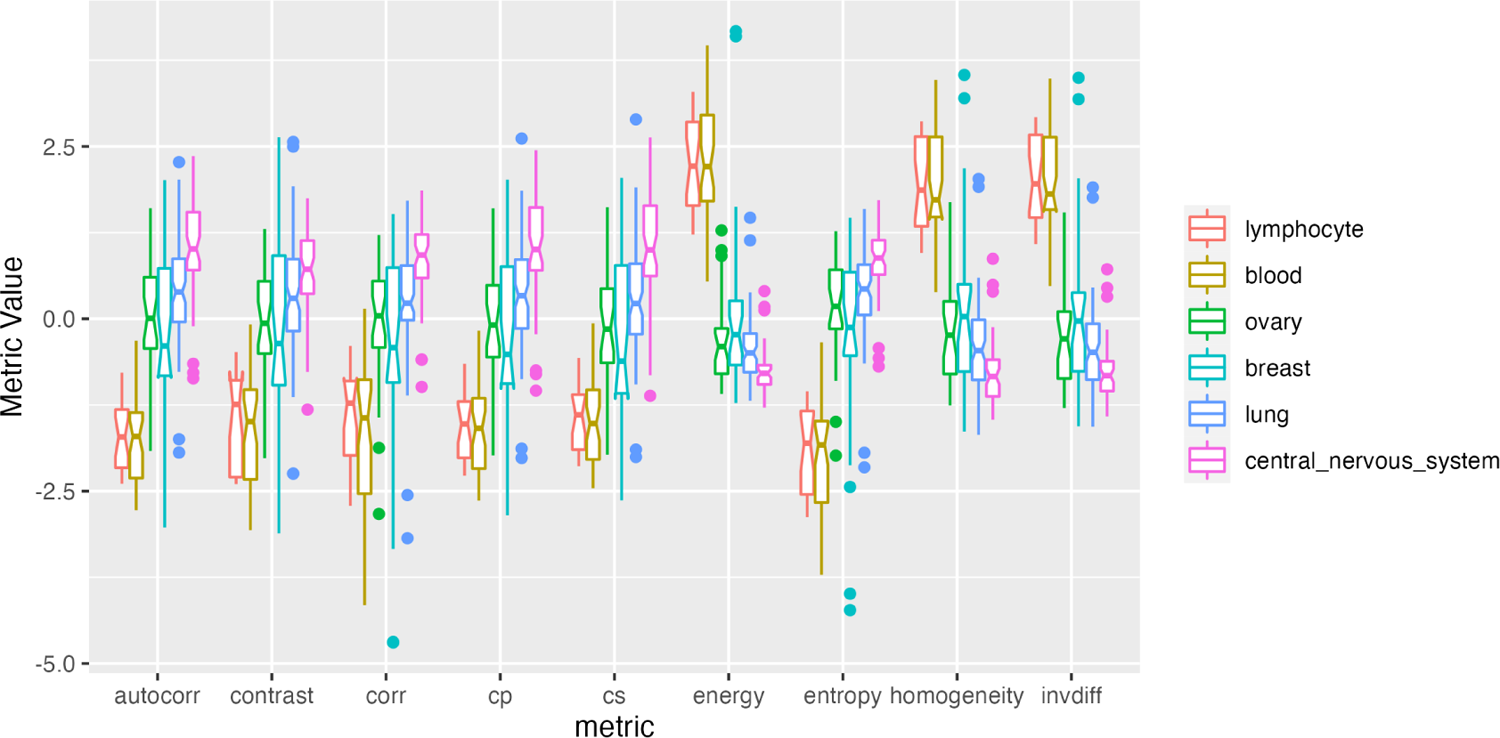
Metrics calculated from gene expression in primary samples of different lineages within the EGFR subnetwork. Gene expression data from the CCLE database was extracted for the genes in the EGFR pathway. Metrics were calculated on this sub-network across 6 of the most represented cancer lineage types in the dataset. Epithelial tumors are separated from non-epithelial tumors in the dataset.

**Figure S4.**
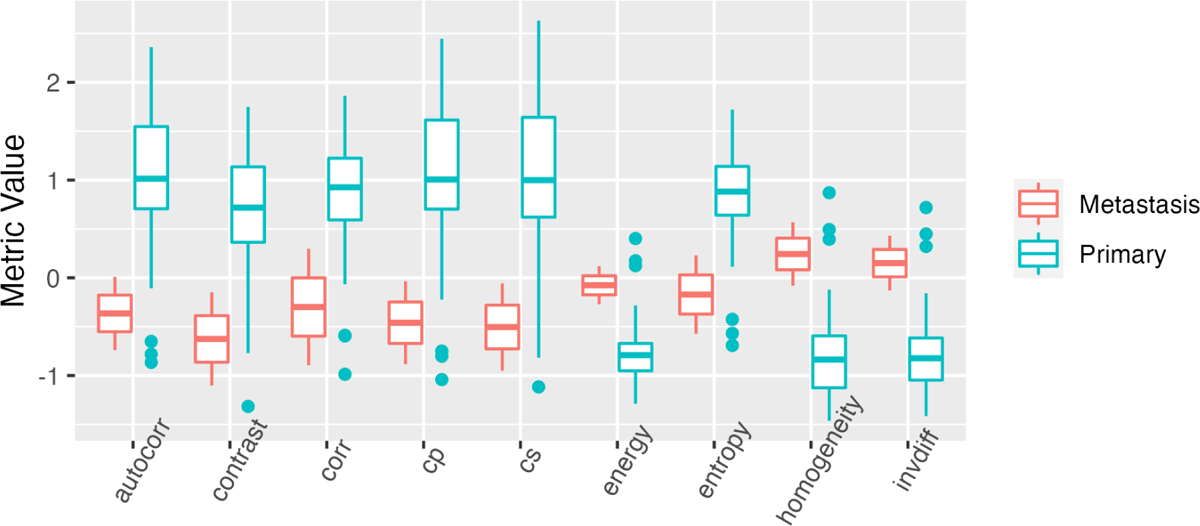
Primary vs Metastatic central nervous system samples for EGFR expression subnetwork using expression values from the CCLE database

---

**Table S1.** Statistical notation for quantities required in feature calculations.

| Definition | Term |
| --- | --- |
| Probability neighboring nodes have weights $(i, j)$ | $p(i, j)$ |
| Marginal probability distribution over rows | $p_x(i, j)$ |
| Marginal probability distribution over columns | $p_y(i, j)$ |
| mean of $p_x(i, j)$ | $\mu_x$ |
| mean of $p_y(i, j)$ | $\mu_y$ |

